# Extreme Genomic Makeover: Evolutionary History of Maternally-transmitted Clam Symbionts

**DOI:** 10.1101/2020.02.01.930370

**Authors:** M Perez, C Breusing, B Angers, YJ Won, CR Young

**Affiliations:** Department of Biological Sciences, Université de Montréal, Montreal, Canada; Graduate School of Oceanography, University of Rhode Island, Narragansett, USA; Division of EcoScience, Ewha Womans University, Seoul, South Korea; National Oceanography Centre, Southampton, UK

## Abstract

Given their recent switch to a vertically-transmitted intracellular lifestyle, the chemosynthetic bacteria associated with deep-sea vesicomyid clams are an excellent model system to study the processes underlying reductive genome evolution. In this study, we provide the first estimates of the relative contributions of drift, recombination and selection in shaping the ongoing reductive genome evolution in these symbionts. To do so, we compared the genomes of endosymbionts associated with 11 vesicomyid clam species to that of closely related free-living bacteria and their respective hosts’ mitochondria. Our investigation confirmed that neutral evolutionary processes were the dominant driver of reductive genome evolution in this group and highlighted the important role of horizontal gene transfer in mitigating genome erosion. Finally, a genome-wide screen for episodic positive selection across the symbiont phylogeny revealed the pervasive role of selective processes in maintaining symbiont functional integrity.

## Introduction

The evolution of biological complexity includes many examples of symbiotic associations. For example, the early evolution of the eukaryotic cell involved multiple endosymbiotic events leading to mitochondria and plastids ^1,2^. More recent examples include associations of metazoans with intracellular bacteria^3–6^, including the well-studied associations of insects and *Buchnera* proteobacterial symbionts ^7^. These associations have profound consequences for both host and symbiont, ranging from alterations of sex-ratio in insect hosts to providing nutrients that are otherwise unavailable in the host’s habitat. Some intracellular symbionts are transmitted from parent to offspring of hosts through the germline (i.e. vertical transmission), while others are acquired from the environment every generation ^6^. The mode of transmission strongly affects the evolution of the microbial partner in these symbioses, as the genomes of vertically transmitted symbionts all seem to follow the same process of reductive genome evolution (RGE) regardless of their phylogenetic origin, host, or habitat. Compared to their free-living counterparts, the genomes of host-restricted symbionts are smaller, contain fewer genes, and are enriched in AT ^8,9^. A prime example is the genomes of cellular organelles such as mitochondria and plastids which are extremely streamlined compared to their bacterial cousins ^10^. Symbiont genome evolution is thought to follow two main stages ^11^. Following host restriction, symbionts undergo rapid genome erosion as they lose non-essential genes through pseudogenization and deletions ^12,13^. Then, symbionts enter a “stabilizing phase”. At this point, their genomes are streamlined, redundant genes and functions are lost ^14^, and the effective rate of deletion diminishes ^15^. This process might be largely neutral due to the reduced effective population size of host-restricted taxa.

The pea aphid/*Buchnera* symbiosis and several other insect/bacteria models support the neutral hypothesis. Captured symbionts experience successive bottleneck events during their transmission that reduce their effective population size and increase genetic clonality. As a consequence, genetic drift increases relative to selection in these taxa ^16–18^. Under these circumstances, elevated mutation load (i.e. the Muller’s ratchet ^19^) and genetic erosion might lead to the functional death of the symbiont lineage ^17,20–22^ unless compensating mechanisms such as gene transfer to the host nucleus or compensatory mutations alleviate the genetic load. Likewise, deep-sea taxa exhibit evidence of nearly neutral processes affecting evolutionary rates due to reduced population sizes in vertically transmitted symbionts ^23^. Other metazoan/microbial symbioses highlight the importance of selection in shaping reductive genome evolution. For instance, symbiont traits that are beneficial for the host are likely to experience increased selective pressures, while selection may be relaxed on genes that are functionally redundant ^8^. Red Queen dynamics are expected to occur in obligate symbioses to maintain the host-symbiont specificity and the functioning of cyto-nuclear interactions through speciation ^20^. Unfortunately, the role of positive selection has often been ignored in studies of symbiont genome evolution and broad screens for positive selection have almost never been performed.

The intracellular sulfur-oxidizing bacteria associated with deep-sea vesicomyid clams (Bivalvia: Vesicomyidae: Pliocardiinae) represent an ideal model to address the neutral and selective processes driving reductive genome evolution. The symbionts are found within the epithelial cells of their host’s gills and provide them with chemosynthetically derived food. They are vertically transmitted to the next generation through the eggs ^24,25^ and generally show co-speciation with their hosts ^26,27^. It is assumed that symbiont capture in these animals was a single event that, based on fossil and molecular information, happened before their radiation about 45 Mya ^28^, an acquisition that is much more recent than that of other well-studied models such as the aphid/*Buchnera* (∼ 200 Mya ^29^) and nematode/*Wolbachia* (∼100Mya ^30^) symbioses. Today, the hosts represent the most diverse group of deep-sea bivalves ^31^, with 173 described species present in a variety of reducing habitats worldwide from hydrocarbon seeps on continental margins to hydrothermal vents on mid-ocean ridges ^32–34^. A comparative study of the first two sequenced vesicomyid symbiont genomes ^35,36^ indicated that they possessed intermediate genome sizes and level of AT enrichment compared to other host-restricted symbionts ^11^. The symbionts of deep-sea vesicomyid clams group into two divergent clades: Clade I (associated with hosts of the *gigas* group), and Clade II (associated with all other lineages of vesicomyid hosts) ^37^. The genomic characteristics of Clade I symbionts indicate that this group is in an advanced state of reductive genome evolution compared to Clade II. However, in contrast to the well-studied pea aphid/*Buchnera* association, which has been in a state of stasis for 50 Myrs ^38^, the evolutionary processes responsible for remodeling the genomes of vertically transmitted symbionts appear to be still operating in the vesicomyid clam symbiosis. Conspicuous bottlenecks during transmission ^25^ and loss of DNA repair genes in several lineages ^37^ suggest that neutral processes and mutational pressures are driving RGE in vesicomyid symbionts, although this hypothesis has not been formally tested.

In this study, we aim to assess the relative contribution of neutral and selective processes to genome evolution in the maternally transmitted symbionts of deep-sea vesicomyid clams. Specifically, we test the hypotheses that genetic drift is the main driver of RGE in these symbionts and that diversifying selection has shaped their genome to maintain host-symbiont epistasis throughout the evolutionary history of the symbiosis. To do so, we applied comparative methods to the symbiont genomes of 11 vesicomyid deep-sea clam taxa representative of the diversity of Clade I and Clade II, the mitochondrial genomes of their respective hosts, and two of their close free-living relatives: the environmentally acquired gill symbiont of the hydrothermal vent mussel *Bathymodiolus thermophilus* and the free-living bacteria of the SUP05 group, which are marine chemoautotrophic Gammaproteobacteria found in hypoxic waters ^39,40^.

## Results

### Host mitochondrial and symbiont phylogenies

Host mitochondrial genomes from the lineages examined in this study possess identical gene orders and contents as previously published mitochondrial genomes ^41,42^. The phylogeny constructed with mitochondrial genome data (Figure 1A) is congruent with the known host phylogenetic relationships based on multilocus sequence data and the *COI* phylogeny ^31^. Structural variation is, however, present. We observe the previously described noncoding structural variation, hypothesized to be the control region, between the *tRNA*^*Trp*^ or *tRNA*^*His_2*^ and *ND6* loci ^41–43^ but we were unable to resolve this region with the current sequence data. We also found the *COX2* gene varies in length among taxa (range: 1005-1452bp). All protein-coding genes in the mitochondrial genomes were screened for selection using the adaptive branch-site random effects likelihood method. Interestingly, the *COX2* gene exhibited evidence for episodic diversifying selection on multiple branches of the phylogeny.

**Figure 1.**
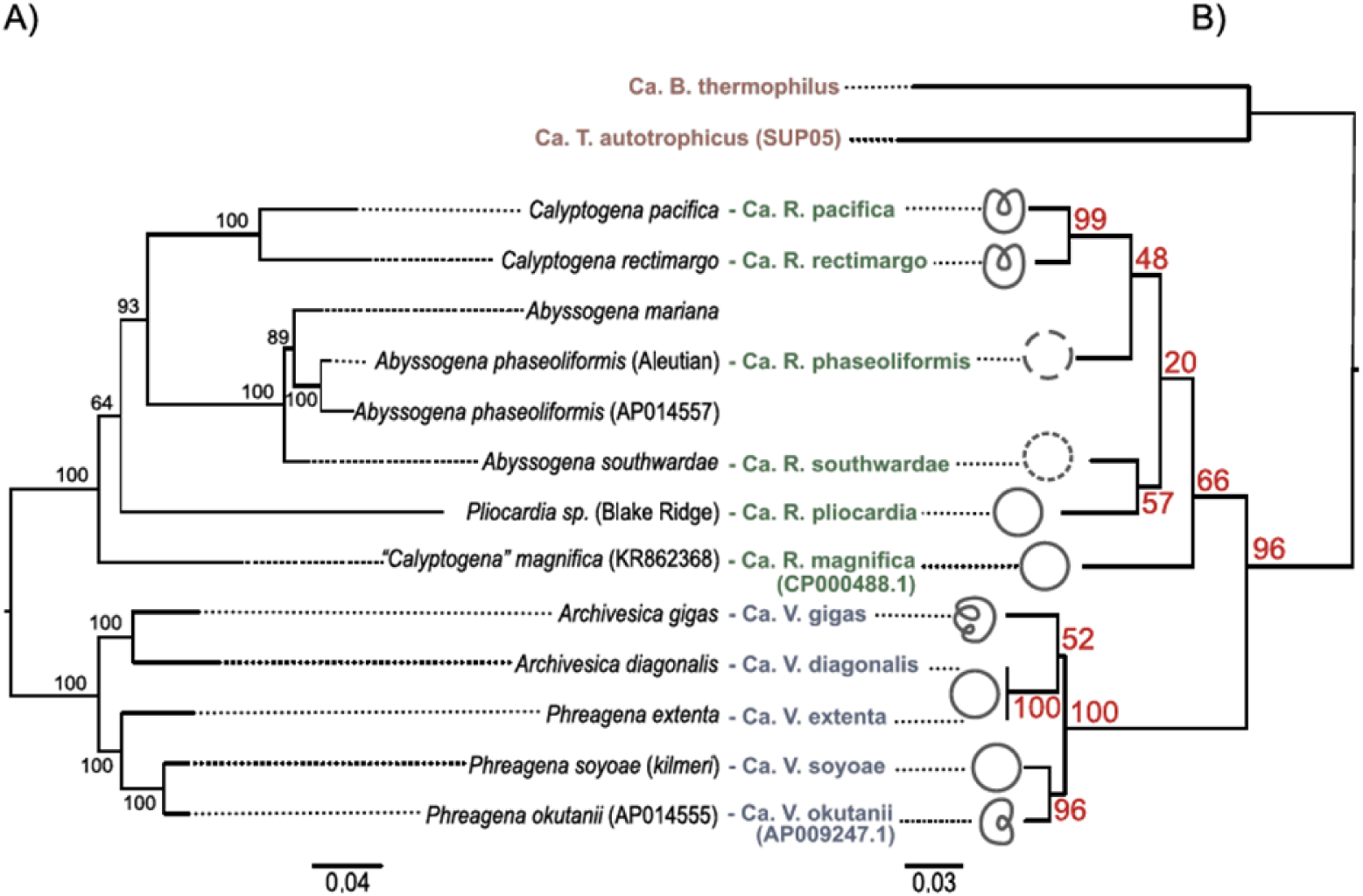
Host and symbiont phylogenomic estimates. A) Neighbor-joining phylogeny based on genetic distance (GTR model) between genome-wide alignments of mitochondrial genomes (15272 bp). Numbers in black are bootstrap values. B) Neighbor-joining phylogeny based on genetic distance (GTR model) between genome-wide alignments of symbiont (Clade I; *Ca*. Vesicomyosocius in blue, Clade II; *Ca*. Ruthia in green) and free-living (in red) genomes (761866 bp). Chromosome schemes showing genome inversions and assembly fragmentation are displayed at the end of the branches. Refer to text for a description of the genome structures. Numbers in red are the genome-wide mean covariance factors; they represent the percentage of protein-coding genes supporting each split of the phylogeny.

Genome size and GC content for the 11 symbiont assemblies in our study varied from 1.02Mb to 1.59 Mb and 31% to 37% GC, respectively (Table 1). The number CDS ranged from 939 in *Ca*. V okutanii to 2210 in *Ca*. R. phaseoliformis. Following initial nomenclature, the symbiont lineages are referred to by the previously erected genera for this group, *Candidatus* Vesicomyosocius for Clade I, and *Candidatus* Ruthia for Clade II symbionts, followed by host species names ^35,36,44^. This classification at the genus level is coherent with both the phylogenetic definition based on 16S identity (inter-genus identity < 95% ^45^) and functional definition based on criteria of genetic isolation ^46^ (see Symbiont genome structure and recombination)

**Table 1.**
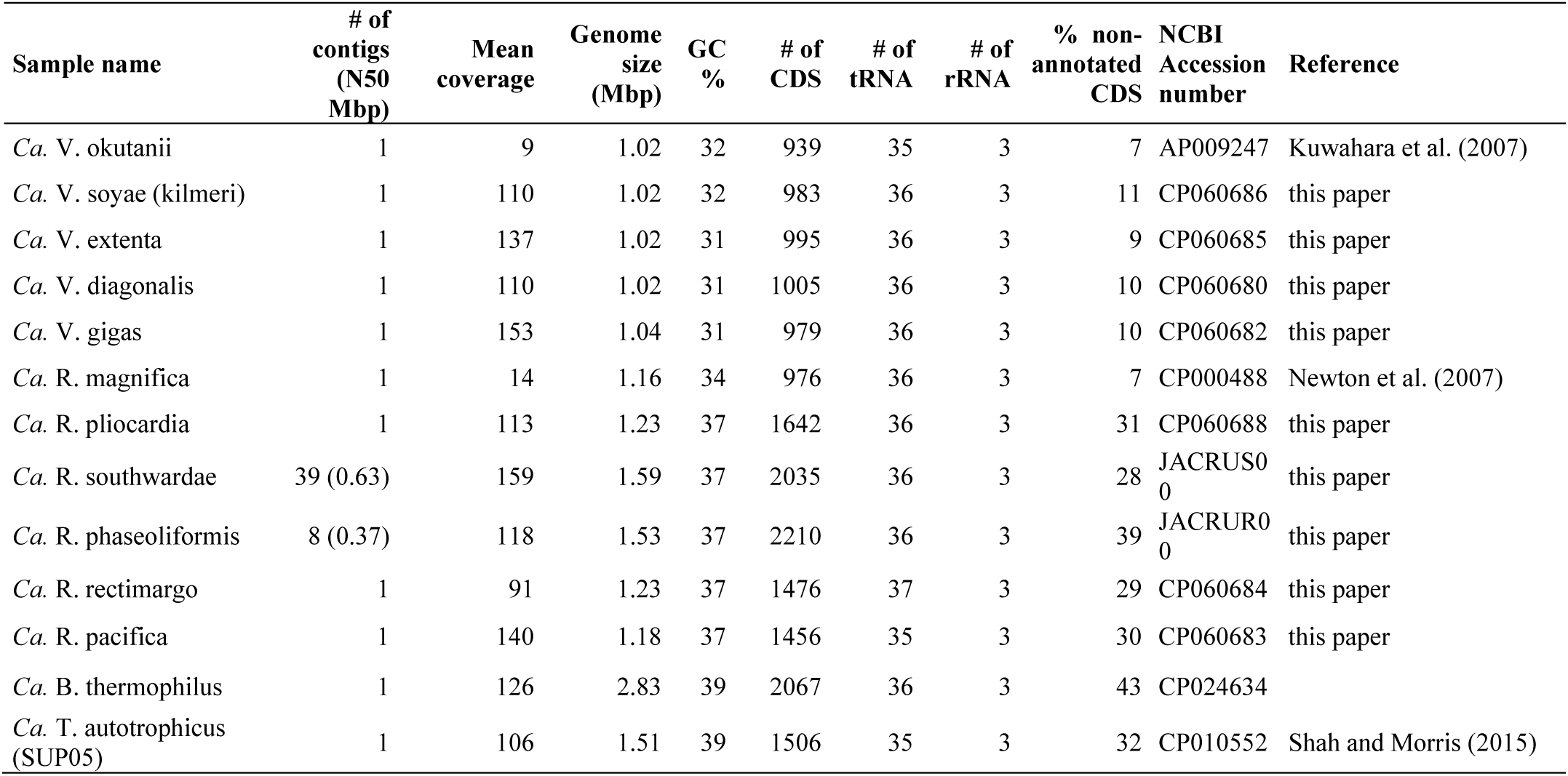
Annotation statistics for symbiont and free-living genomes in this study.

Examination of the mitochondrial and symbiont phylogenies (Figure 1) shows good concordance for all lineages except one. The symbiont lineages of *Ca*. V. diagonalis and *Ca*. V. extenta are nearly identical whereas their respective host mitochondrial lineages are divergent. The donor lineage in this recent symbiont replacement appears to be *A. diagonalis*. It is noteworthy that these clams were both collected from sites in Monterey Canyon. Pairwise comparison of mitochondrial and symbiont genome-wide synonymous divergence indicates faster evolutionary rates in the mitochondria compared to the symbionts in almost every holobiont pair (Figure 2). Within the symbionts, we detect signatures of elevated substitution rates on the branch leading to Clade I: the symbiont pairs across the Clade I-Clade II bipartition have significantly higher divergence than the others even when controlled for host divergence (1 < dS_mito_<2).

**Figure 2.**
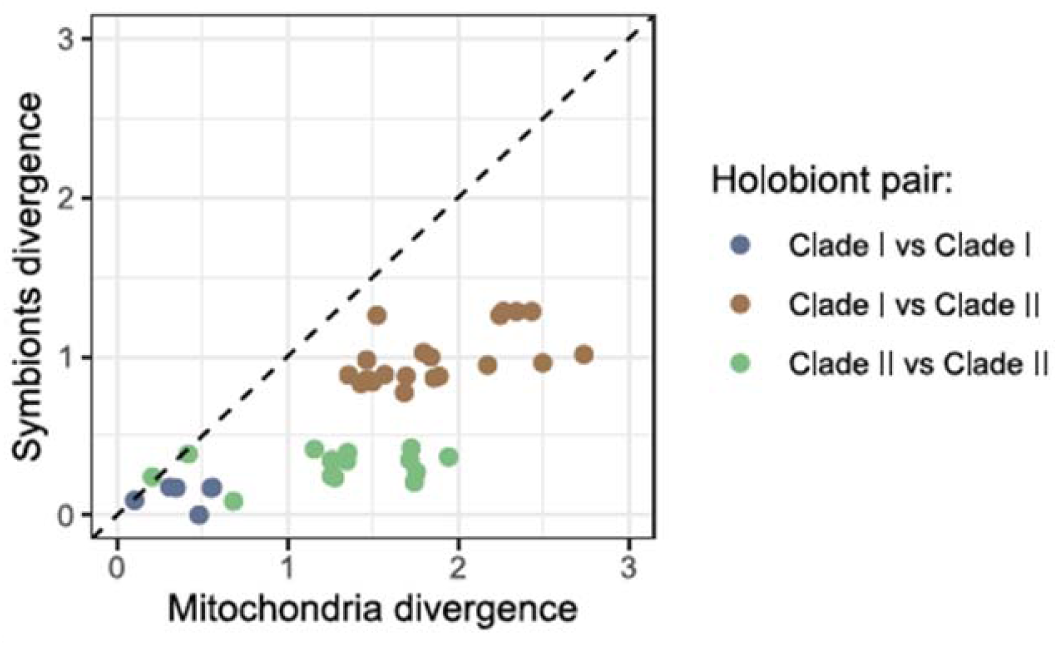
Relationship between symbiont and mitochondrial divergence. For each holobiont pair, host and symbiont divergences are expressed as pairwise synonymous substitutions rates (dS) in their respective genomes.

### Symbiont genome structure and recombination

Free living bacteria associated with *B. thermophilus* and *Ca*. T. autotrophicus shared about 1 Mbp of their genomes with the clam symbionts. Permutation analysis of locally collinear blocks (i.e. long fragments of aligned genomes) with GRIMM (http://grimm.ucsd.edu/cgi-bin/grimm.cgi) showed that at least 18 inversion events occurred between the genome of the *B. thermophilus* symbiont and that of the *Ca*. R. magnifica reference. Fewer rearrangements (3 inversions) were observed between SUP05 and *Ca*. R. magnifica.

Genome structure among the clam symbionts was also variable (Figure 1B). The previously reported *Ca*. V. okutanii genome ^36^ possesses one inversion compared to that of *Ca*. R. magnifica ^35^ but that of *Ca*. V. okutanii’s closest relative, *Ca*. V. soyoae, does not. The genomes of *Ca*. R. pacifica and *Ca*. R. rectimargo share a single inversion distinct to that of *Ca*. V. okutanii. Two other inversions were found in the *Ca*. V. gigas genome. Finally, read-mapping to the consensus assemblies for *Ca*. R. phaseoliformis and *Ca*. R. southwardae suggested the presence of intra-host structural variation in these symbionts.

Applying Bayesian concordance analysis to all core protein-coding genes, we detect a large amount of recombination among symbiont lineages, though recombination is not randomly distributed. We observe no recombination between members of Clade I and II, but recombination is occurring within these genera (Figure 1B). Strikingly, much less topological concordance was found in Clade II – more than 40 different topologies were necessary to fully represent the diversity of conflicting phylogenetic signals – compared to that of Clade I whose phylogeny was fully represented by 5 different trees. Within Clade I, conflict originates from the uncertainty of the position of *Ca*. V. gigas. Only 50% of the genes support its position in the phylogenetic tree issued from the concatenated core genome alignment (Figure 1B). Other well supported positions for this species are at the base of the clade (supported by 27% of genes) and closer to the group composed of *Ca*. V. soyoae and *Ca*. V. okutanii (supported by 20% of genes). Within Clade II, only the grouping of the sister species *Ca*. R. rectimargo and *Ca*. R. pacifica is supported by the topologies of all genes while the positions of other species have low support.

### Gene conservation across symbionts and free-living bacteria

#### Genes of free-living and horizontally-transmitted bacteria missing in vesicomyid symbionts

The genomes of the free-living bacteria contained many large (> 5kb) contiguous sections that were not found in the symbionts. These genomic islands were mostly composed of unannotated genes and mobile elements (transposases, integrases, prophage genes) (Table S1). We found more selfish genetic elements in the genome of the *Bathymodiolus* symbiont than in that of SUP05. The genomic islands found in the two genomes also encoded several gene clusters of particular functional interest described below.

Unsurprisingly for a bacterium living in a metal-rich hydrothermal environment, the *B. thermophilus* symbiont genome possesses genes for resistance against heavy-metal toxicity such as a multi-copper oxidase (*mmcO*), a copper ion exporting ATPase (*copB*), cobalt-zinc-cadmium resistance proteins (*czcD* and c*zcCBA*), and a chromate transport protein (*chrA*). The genomic islands of the mussel symbiont also carried full operons for three different defense systems; a type I restriction and modification system (*hsdRMS*), a CRISPR-Cas type II system (*cas9, cas1, cas2, cas4*), and a type II toxin-antitoxin system (*vapCB*). Finally, this genome possesses a 23kb hydrogenase operon that has 83% and 82% identity to that of the symbionts of *Bathymodiolus septemdierum* ^47^ and *B. puteoserpentis* ^48^, respectively. The representative SUP05 genome contained a 21kb motility locus, comprising a type IV pilus biogenesis operon (*pilA, pilB, pilC, pilT, pilQ, pilY1*), and a toxin-antitoxin locus (*higAB*), that was not found in the other genomes. Furthermore, this genome possessed two additional smaller genomic islands (6kb and 15 kb) encoding a nitric oxide reductase (*norCBQD*), and a periplasmic nitrate reductases (*napAB*), respectively, which clustered with sulfur covalently binding protein genes (*soxYZ*).

#### Gene content in vesicomyid symbionts

The symbionts of Clade I and Clade II possessed essentially a subset of the genes found in the free-living lineages. Indeed, sequence-based comparisons of free-living lineages to the symbionts revealed that many genes present only within the symbiont lineages are hypothetical genes with unknown function resulting from the degeneration of ancestral genes, as indicated by premature stop codons, frameshifts, and loss of neighboring genes (Table S1). These pseudogenes were more prevalent in the genomes of Clade I than Clade II symbionts. In many instances, homologous regions within the Clade I symbiont genomes were instead characterized by large deletions. In general, patterns of gene decay were more variable within Clade II than Clade I. Genes were overall more conserved within *Ca*. R. southwardae, *Ca*. R. phaseoliformis and *Ca*. R. pliocardia than in other lineages. Among the *Ca*. Ruthia symbionts, gene degeneration was most pronounced in *Ca*. R. magnifica, which possessed a conservation pattern closer to that of the Clade I lineages (Figure S1B).

### Genome-wide pattern of relaxed selection

Codon usage bias was reduced in the symbiont lineages compared to their free-living relatives. Furthermore, symbionts in Clade I showed reduced bias and variance compared to Clade II (Figure 3A). The CDC values of core protein-coding genes were significantly correlated between lineage pairs both at the clade and species level (Pearson’s test p-value <0.001; Figure 3B, Table S2), suggesting that the reduction in codon usage bias in the vertically transmitted symbionts result from a genome-wide reduction of the efficacy of purifying selection.

**Figure 3.**
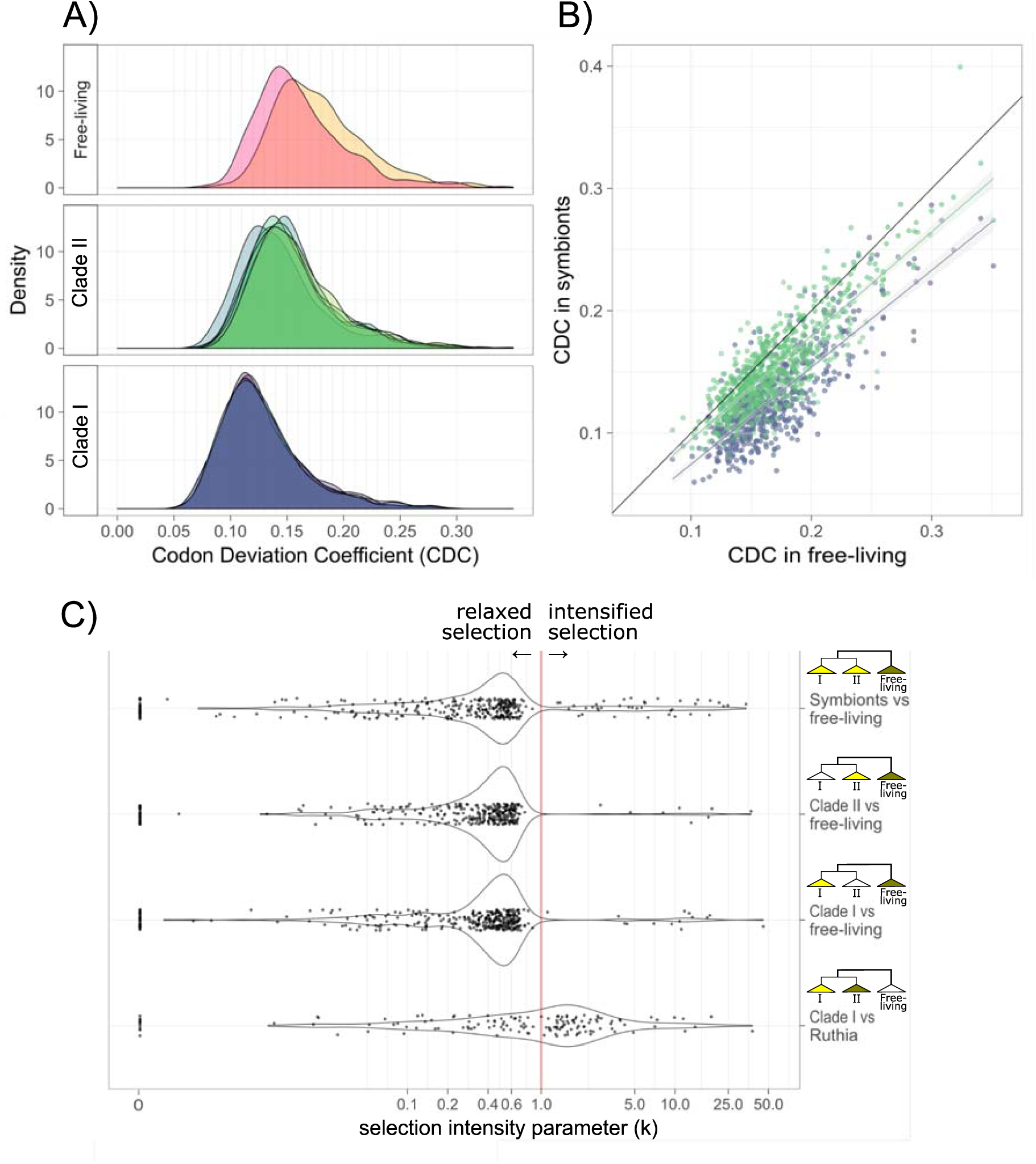
Codon bias in symbionts and free-living. A) Codon Deviation Coefficient (CDC) spectra for each genome (all protein-coding genes). Within the free-living, yellow: *Ca*. B. thermophilus; red: *Ca*. T. autotrophicus. B) Correlation between the average CDC of free-living, *Ca*. Ruthia (green) and *Ca*. Vesicomyosocius (blue) core genes. Linear regressions are shown. CDC values vary from 0 (no bias) to 1 (maximum bias). C) Selection parameter (k) spectra of genes for which a significant change in selection was detected by RELAX. Note that k is on a log scale.

RELAX analysis revealed intensified selection in the vesicomyid symbionts compared to free-living bacteria for less than 5% of the core orthologous genes, while relaxed selection was detected in more than half of the core gene set (Figure 3C, Table S3). The magnitude of relaxation (k<1) was in the range of that observed in insect endosymbionts ^49^ but was not correlated to codon bias. Genes exhibiting intensified and relaxed selection represented a multitude of metabolic functions, but genes under relaxed selection were enriched in the protein metabolism, nucleoside and nucleotides, and DNA metabolism categories while genes under intensifying selection were more likely to be associated with respiration, cell wall and capsule, and sulfur metabolism. However, we did not find increased relaxation in the symbionts of Clade I compared to Clade II. Indeed, fewer genes exhibited significant change in selection pressure (intensified or relaxed) between these groups than between symbionts and free-living bacteria, and about the same proportion of genes under relaxed and intensified selection was found in both clades.

### Genome-wide screen for positive selection

The symbiont genes that passed the inclusion criteria to be screened for selection (see methods) included 652 loci. The application of the adaptive BS-REL method yielded 223 genes with significant evidence for episodic diversifying selection along branches in the phylogeny. Selection is distributed throughout the evolutionary history of the group (Figure S1A, and Table S4) with most selection occurring on the branches discriminating free-living bacteria, Clade I, and Clade II (branches a, b, and c in Figure 4), as well as within the *B. thermophilus* symbiont and SUP05 lineage (43 and 37 genes, respectively). Eighty-five percent of the loci that exhibited unequivocal evidence of selection was assigned to SEED categories (Figure 4, Table S5). Within each clade and along each of the main branches, these selected loci were not equally represented amongst cellular functions of the core genome (hypergeometric tests p-values < 0.001). Genes in overrepresented functional categories are presented in Table 2. The complete list of selected genes is available in Table S4.

**Table 2.**
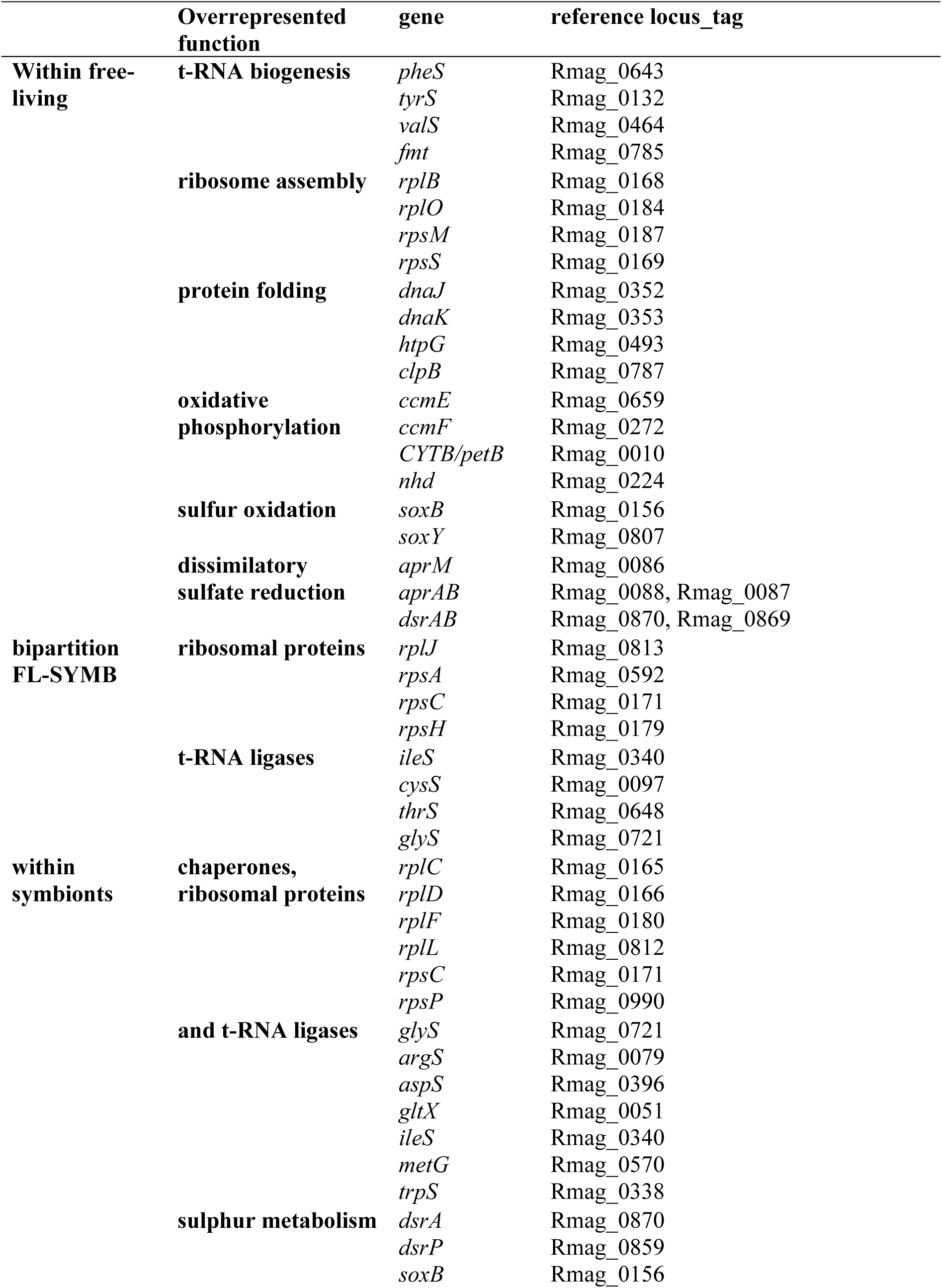
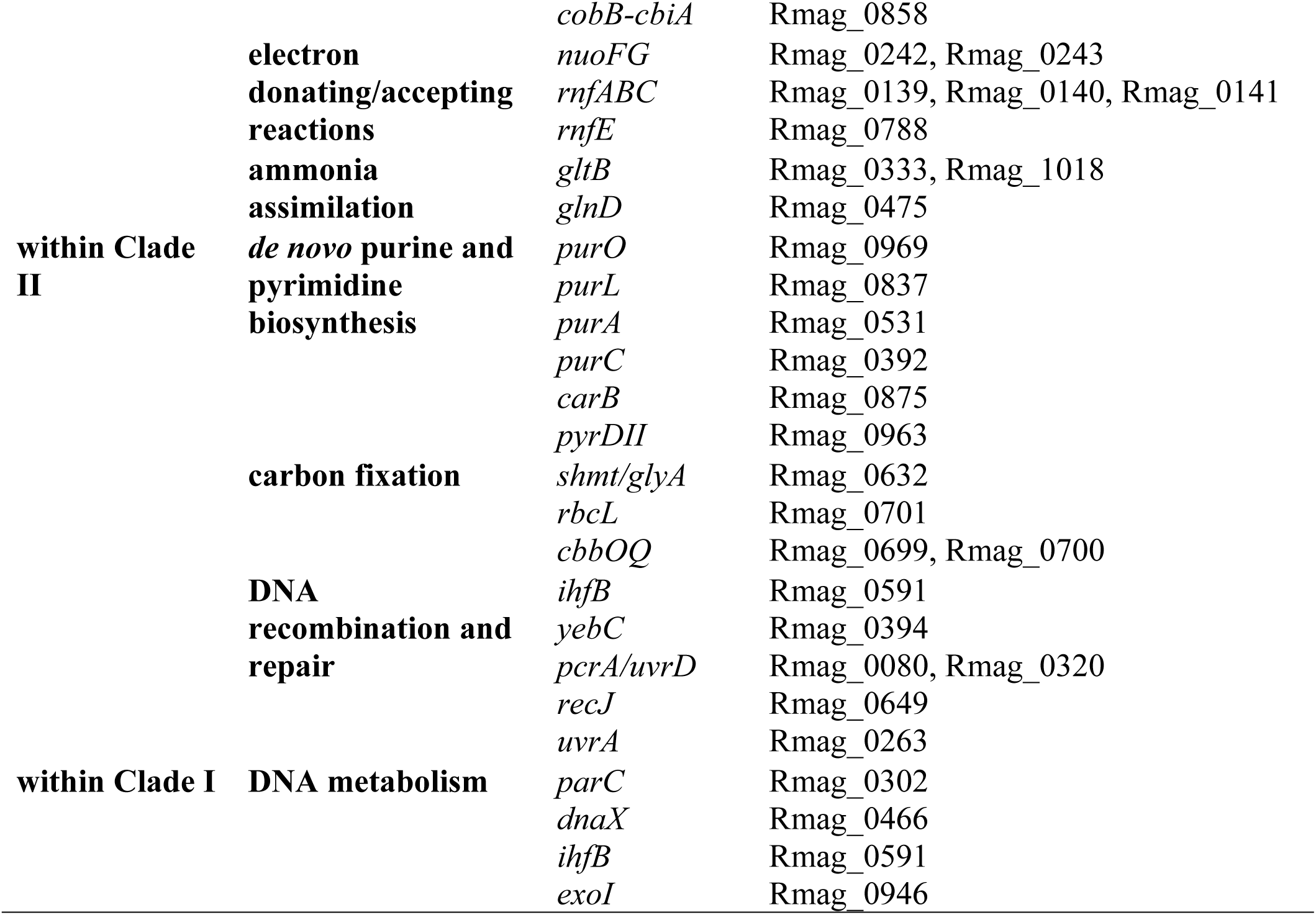
Overrepresented functional categories for genes exhibiting significant evidence for episodic diversifying selection

**Figure 4.**
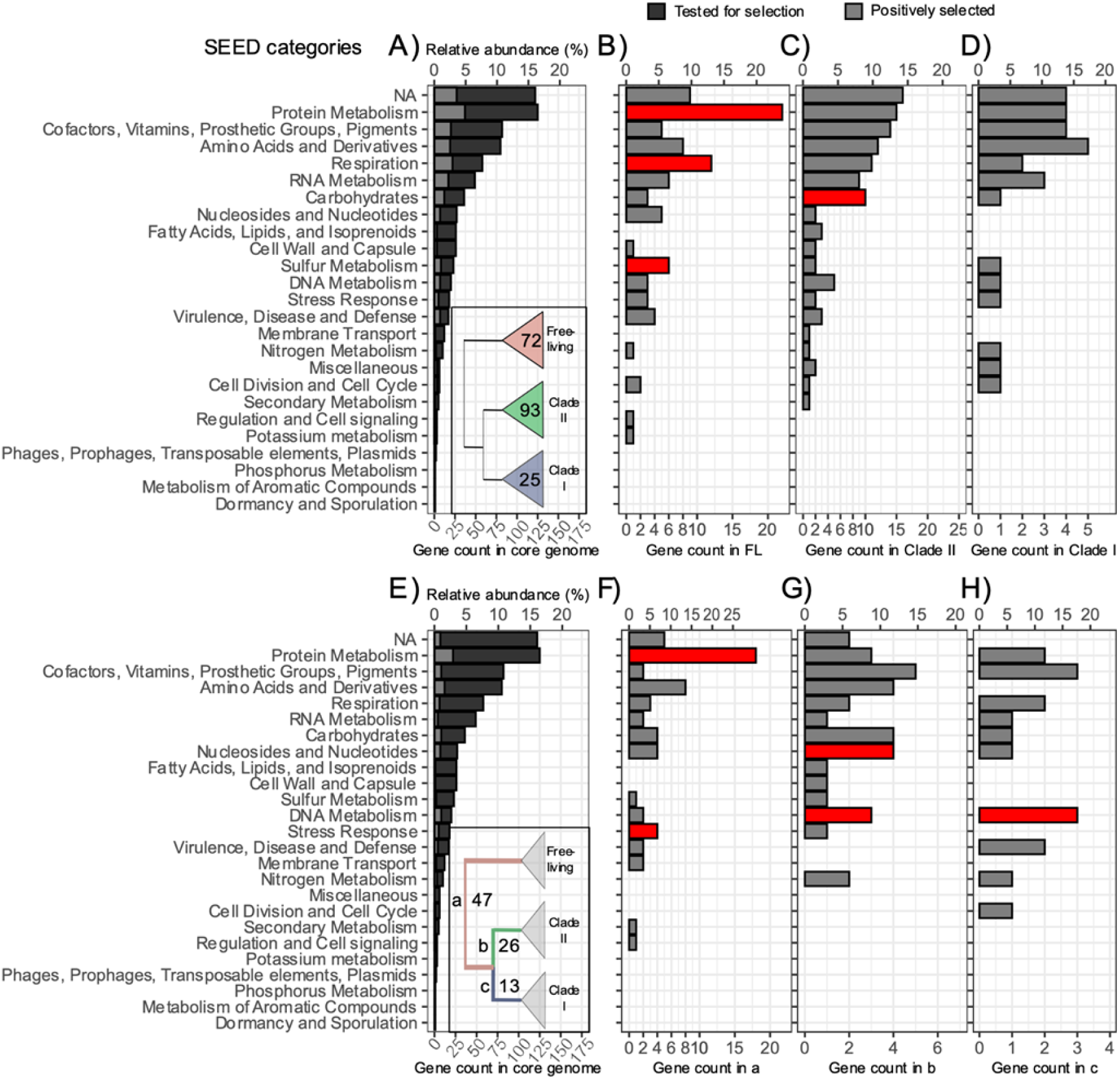
SEED category distribution of core genes under episodic diversifying selection within phylogenetic clades (A, B, C, D), and on partitioning branches (C, D, E, F). A) Distribution of all non-recombining core genes (dark grey, 652 loci) and loci under selection within the free-living, *Ca*. Ruthia, and *Ca*. Vesicomyosocius clades (light grey, 168 loci). The number of loci selected within each clade is represented in the inset. B) Genes under selection within the free-living. C) Genes under selection within *Ca*. Ruthia. D) Genes under selection within the *Ca*. Vesicomyosocius. E) Distribution of all non-recombining core genes (dark grey, 652 loci) and loci under selection on all partitioning branches (light grey, 80 loci). The number of loci selected on each branch is represented in the inset. F) Genes under selection on branch a. G) Genes under selection on branch b. H) Genes under selection on branch c. Note that genes may be represented in multiple functional categories and multiple clades or branches. SEED categories significantly overrepresented (in red) and underrepresented (in blue) in the groups compared to the core genome are highlighted. Refer to text for further breakdown of these categories. NA: no functional annotation.

#### Selection within free-living bacteria

Amongst the free-living lineages, a larger than expected number of genes associated with protein metabolism, respiration, and sulfur metabolism were under selection (Fisher tests p-value <0.05). These included genes involved in ribosome assembly, t-RNA biogenesis, protein folding, oxidative phosphorylation, sulfur oxidation, and dissimilatory sulfate reduction. On the bipartition between the free-living and symbiont groups, additional genes associated with protein metabolism were positively selected, including ribosomal protein genes, and the t-RNA ligase genes.

#### Selection within symbionts

Many genes coding for chaperones, ribosomal proteins, and t-RNA ligases were under selection within the symbiont phylogeny. In addition, we found evidence for selection in metabolic genes that are central to the chemosynthetic role of these symbionts. Several genes involved in sulphur metabolism (i.e. *dsrA, dsrP, soxB, cobB-cbiA/dsrN*) and electron donating/accepting reactions were under selection. Two genes involved in ammonia assimilation (*gltB*, and *glnD*) also exhibited evidence of selection within both symbiont clades. Within Clade II and along the branch partitioning this group, there was an over-representation of selected genes involved in *de novo* purine and pyrimidine biosynthesis, carbon fixation, and DNA recombination and repair. Selection within Clade I favored additional genes broadly associated with DNA metabolism. Notably, 60 genes showed evidence for positive selection in multiple branches of the phylogeny, including 44 genes within the symbiont phylogeny. These genes were mostly associated with protein metabolism.

## Discussion

### Reductive genome evolution is still ongoing in the clam symbionts and is driven by neutral processes

Comparative analyses of the first two reference genomes of vesicomyid clam endosymbionts revealed variation in genome structure, genome characteristics, and genome composition between distantly-related symbiont species ^11^ suggesting that RGE might still be ongoing in this group. Our results confirm these early findings and reveal additional genomic variation among the deeply diverging lineages. These findings expand the ranges of genome size, genome content and GC% considerably.

As in other models of recently acquired bacteria ^22,50^, gene content differed greatly between vesicomyid symbiont genomes indicating that the different lineages are independently losing genes. The presence of structural variation and putative pseudogenes (Figure S2) within the vesicomyid symbiont genomes suggest that these symbionts have not yet reached a stable streamlined state as those of the *Buchnera* or *Paulinella* symbionts ^15,38^. Comparing the clam symbionts to their free-living relatives revealed reduced GC%, a reduction in codon usage bias, pseudogenization, and evidence for reduced purifying selection in the vast majority of genes. Taken together, these observations support the nearly neutral theory of RGE, driven by a reduction of effective population size in these taxa.

Finally, in agreement with the findings of Stewart et al. ^27,51^, Decker et al. ^52^, and Ozawa et al. ^53^, we detected no recombination between Clade I and II symbionts even though some of the host taxa co-occur ^54–56^. These findings imply that there is enough molecular and ecological divergence between the two clades for clonal interference and/or strong host-symbiont epistatic interactions to constrain symbiont exchange ^20,52^. Thus, our results support the nomenclature initially put forward by Newton *et al*. ^35^ and Kuwahara *et al*. ^36^ classifying the symbionts from Clade I and II into two distinct bacterial genera, *Ca*. Vesicomyosocius and *Ca*. Ruthia. For clarity, we will keep referring to these two genera as Clade I and Clade II in the rest of the discussion.

### Reductive genome evolution is exacerbated in non-recombining symbionts

Clade I symbionts are in a more advanced state of RGE than the others. Indeed, compared to Clade II, their genomes are smaller and lower in GC%, possess fewer genes and pseudogenes, and exhibit less codon usage bias. The genomes of Clade I symbionts are also more homogeneous. Patterns of gene conservation suggest that much of the loss in this group happened after its speciation but before its radiation, a period of roughly 20Mys ^26,31^. Together with increased substitution rate on its diverging branch these results show that the ancestral Clade I lineage experienced an episodic acceleration of reductive genome evolution. It is likely that the increased level of genome reduction in Clade I results from a reduction of homologous recombination in the ancestor of the group exacerbating Muller’s ratchet ^57^. Drift-driven loss of recombination machinery may have strongly reduced the rate of genetic exchange among the symbionts in this genus. Indeed, essential genes of the RecF and RecBCD pathways for homologous recombination appear to be lost in all of the Clade I symbionts ^37^ and while horizontal transfer of genetic material is widespread among symbionts within Clade II it is almost absent in Clade I.

Strong linkage disequilibrium forces whole genomes to sweep in populations that lack genetic exchange capabilities. Hence, the loss of homologous recombination genes should favor symbiont replacement in cases where the divergence between “native” and foreign symbionts is low (i.e. when the foreign symbionts are not too easily outcompeted by those that have co-evolved with the host). In fact, we find multiple examples of symbiont replacement among symbionts of Clade I. For instance, individual clams of the species *P. extenta* have acquired the symbionts of the sympatric species *A. diagonalis*. Likewise, Breusing et al. ^56^ found a population of *A. gigas* carrying the symbionts of the host species *P. soyoae*. Symbiont replacement occurs in several vertically transmitted symbioses ^58–61^ and is speculated to constitute a mechanism for escaping the evolutionary rabbit hole caused by Muller’s ratchet ^20,58,62^. The present data support this notion, and future population genomic studies could determine the prevalence of symbiont replacement and relative rates of recombination in these taxa on more recent time scales.

Despite the lack of recombining machinery in Clade I, one lineage in this genus, *Ca*. V. gigas, showed evidence for recombination. It is puzzling how recombination might be occurring in this species. Breusing et al. ^56^ recently found evidence of unidirectional introgression from *P. soyoae* into *A. gigas*. This mechanism might enable *A. gigas* symbionts to come into contact with other symbionts. Perhaps the recombination in this species is enabled via host-encoded proteins ^63^. Transfer of symbiont genes to the host nuclear genome is possible and should be investigated in future studies. Indeed, evidence for such transfer was recently found by Ip *et al*. ^44^ who identified *Bathymodiolus* symbiont gene homologs in the genome of the *A. marissinica*.

### Putative ecological and evolutionary consequences of RGE

The Muller’s ratchet has been hypothesized to lead to a progressive loss of fitness in host restricted symbionts ^20^. Sympatric populations of symbionts from Clade I and II represent an excellent model to test this hypothesis because of their contrasting reductive stages. For instance, comparisons of the sulfide physiology of the host species *P. soyoae* and *C. pacifica*, which occupy different micro-niches in the same habitat, reveal that *P. soyoae* individuals have lower sulfide oxidation capacities than those of *C. pacifica* ^55^. This could be the consequence of a less efficient sulfide metabolism in *Ca*. V. soyoae resulting from a more advanced state reductive genome evolution in this species compared to *Ca*. R. pacifica. If RGE in the symbionts can restrict their host’s ecological range, contrasting degrees of RGE may put constraints on the potential for genetic exchange across different holobiont species and even promote speciation ^20^. Future observational and experimental studies could help define the evolutionary constraints imposed by both host and symbiont physiology and clarify the role of reductive genome evolution in niche partitioning and speciation.

### Selective processes in the evolutionary history of the symbionts

Contrasting patterns of gene conservation between the symbionts and their free-living relatives are caused by a shift in selective regime in the host-associated bacteria. Genes enabling bacteria to face the challenges of a free-living environment, such as detoxification, anti-viral defense and inter-species competition, were not conserved in the vesicomyid clam symbionts. Furthermore, different patterns of pseudogenization in Clade I and Clade II likely translate to different physiological adaptations at the level of the holobiont. For example, Breusing et al. [in review] found that the two vesicomyid symbiont clades show enzymatic differences related to sulfide oxidation and nitrate reduction and have contrasting dependencies on nickel and vitamin B12 in accordance with adaptations to different ecological niches. In addition, episodes of diversifying selection on genes associated with respiration, ammonia assimilation, and chemosynthesis might reflect the constraints imposed by the diverse selective pressures of host physiology throughout their radiation and niche expansion.

Selective constrains are expected to affect genes involved in host-symbiont interactions. Interspecific communication between eukaryotes and microbes generally involve molecules with distinct motifs produced by the symbiont (e.g., Nod factors, lipopolysaccharides, or peptidoglycans) that are sensed by special receptor in the host ^64,65^. These molecular pathways must experience reciprocal adaptations to persist through speciation and niche expansion. Diversifying selection acting on genes involved in the mediation of host-symbiont interactions such as lipopolysaccharides and peptidoglycans was observed in divergent clades of *Wolbachia* ^66^ and many facultative endosymbionts ^67^. In a recent study, Chong et al. ^68^ performed a genome-wide screen for selection in the *Buchnera* symbionts from the aphid subfamily Aphidinae. Of the 371 protein-coding genes tested, the authors detected 29 positively selected genes representing a variety of metabolic functions including two outer membrane porins (OmpF and OmpA), which are assumed to be important for host interaction.

Surprisingly, in the clam symbionts, we did not detect selection on proteins associated with host-symbiont interactions but found instead a pervasive pattern of diversifying selection that affected many loci related to housekeeping functions such as DNA and RNA metabolism, transcription and translation. Many ribosomal proteins and chaperones showed evidence for episodic positive selection repeatedly throughout the symbiont phylogeny. These results could indicate that the accumulation of slightly deleterious mutations in the symbiont genomes initiates a selective pressure for compensatory mutations ^69,70^. Evidence for such mutations exist in several organelles and symbiont models ^70–73^. For instance, in insect endosymbionts, positively selected loci of the chaperonin GroEL are suspected to permit better protein binding and allow proper protein folding despite mutations affecting their conformation ^71^. Alternatively, these signatures of selection might be in response to other generalized selection pressures such as differences in host habitat (e.g., depth). However, the host mitochondria do not overall seem to be similarly affected making this alternative less likely. Regardless, the pervasive nature of episodic diversifying selection at the level of amino acids in the symbiont genomes suggests that increased drift due to effective size reduction is not the sole driver of molecular evolution in these taxa.

## Conclusion

The vertically transmitted symbionts of deep-sea vesicomyid clams are an ideal model to study the processes of reductive genome evolution, as they constitute a highly diverse group of host-restricted bacteria with varying degrees of genomic reduction. We show that both neutral and selective processes have played a role in the evolutionary history of these symbiont and that factors affecting their clonality have strongly influenced the rate of genome evolution. While the vesicomyid clams have yet to be successfully bred in aquaria, significant progress has been made towards their cultivation ^74^. Examination of the symbionts at the population-level, both within and across individual hosts, will help to decipher the contributions of host physiology, genetic drift, symbiont fitness, cytonuclear incompatibilities, and horizontal gene transfer to their evolution. Additionally, experimental studies on host-symbiont interactions and holobiont metabolism will shed further light onto the role of these symbionts in the ecological partitioning of their hosts.

## Acknowledgements

The Monterey Bay Aquarium Research Institute kindly provided samples from the Vrijenhoek collection for this study. We thank the ships’ crews and submersible pilots involved in the collections for this study, without whose efforts this work would not have been possible. N. Pratt and A. Baylay contributed to sequencing at the National Oceanography Centre Genomics Facility. This work was supported by NERC National Capability funding. MP acknowledges the support of the National Science and Engineering Research Counsil (NSERC) of Canada’s Alexander Graham Bell graduate scholarship and Michael Smith Foreign Study Supplements. CB’s contribution was supported through a postdoctoral fellowship of the German Research Foundation (BR 5488/1-1) and a grant from the United States National Science Foundation awarded to CB’s mentor Roxanne Beinart at the University of Rhode Island (OCE-1736932). BA acknowledges support from NSERC research grant #238600. The genome of *Bathymodiolus thermophilus* was sequenced as part of a project titled ‘Understanding the deep-sea biosphere on seafloor hydrothermal vents in the Indian Ridge (No. 20170411)’ funded to YJW by the Ministry of Oceans and Fisheries, Korea.

## Materials and Methods

### Sample collection and sequencing

Host taxa examined in this study were chosen from the deepest diverging lineages within the Vesicomyidae that are distributed globally in the northern hemisphere (Figure S3) and are representative of the known host diversity ^31^. Specimens of nine clam species were collected between 1996 and 2004 over eight research expeditions (Table 3, Figure S3). Depths of sampling locations ranged from 650–3550m. Samples were dissected aboard ship and then frozen at -80C or were frozen whole at -80C. DNA was extracted from symbiont bearing gill tissue using the DNeasy Blood & Tissue extraction kit (Qiagen, Hilden, Germany) following the manufacturer’s protocol. Host species identification was initially confirmed by sequencing the host mitochondrial *COI* gene using vesicomyid-specific primers ^28^.

**Table 3.**
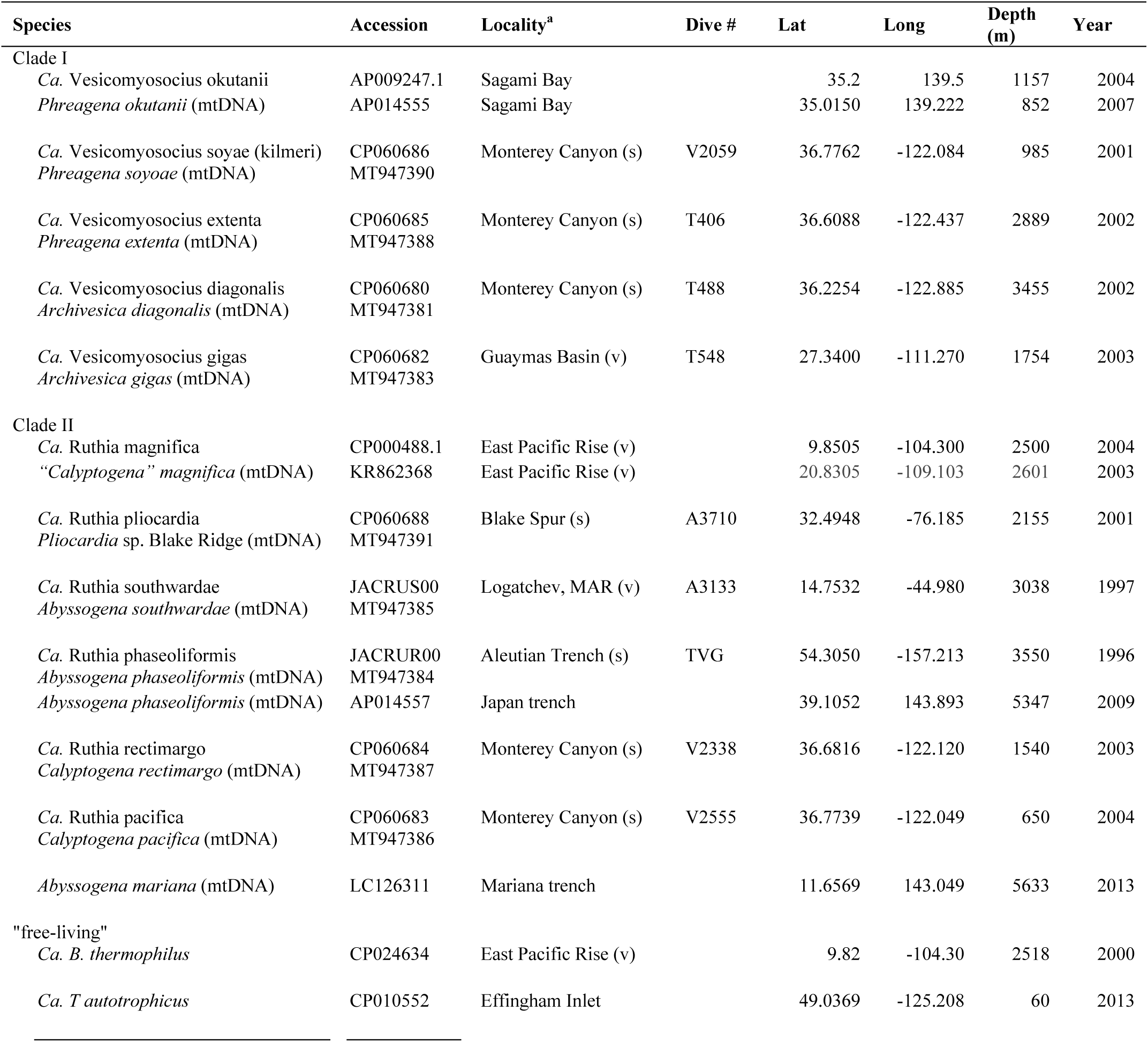
Sampling information and genome accession numbers for taxa in this study

Mixed host and symbiont DNA samples were sequenced in-house on a MiSeq instrument. Genomic DNA libraries were prepared using the KAPA Hyperplus Library Preparation kit (KAPA Biosystems, Wilmington, MA, US) according to kit instructions. Read quality of genomic data was assessed using FastQC ^38^.

### Mitochondrial and symbiont genome reconstruction and annotation

Initial symbiont and mitochondrial assemblies were constructed from the same metagenomic libraries (Table 3) using Velvet ^76^, manually optimizing for *k*-mer size distribution and read depth. Some assemblies were also constructed using the read mapping and assembly functions in Geneious version 10.1.3 ^77^.

Scaffolding and circularization of the symbiont genomes were performed by mapping, extracting and reassembling reads mapping to the extremities of contigs using Bowtie2 ^78^, Samtools ^79^ and SPAdes ^80^, respectively. Mitochondrial genomes were assembled de novo with MITObim ^81^ using as seed a set of initial contigs constructed using the read mapping and assembly functions in Geneious version 10.1.3 ^77^. Mitochondrial genome annotations were produced by the GeSeq application ^82^ using ARWEN v1.2.3 for tRNA prediction, and manually curated with the aid of previously annotated mitochondrial genomes ^41,42^ in Geneious. Mitogenome assembly statistics are presented in Table S6. The symbiont genomes were annotated in RAST ^83^.

### Structural variation and phylogenomic analyses

Host mitochondrial and symbiont genomes were aligned with Progressive Mauve ^84^. Progressive Mauve and GRIMM (http://grimm.ucsd.edu/cgi-bin/grimm.cgi ^85^) were used to identify large-scale structural differences among genomes. Locally collinear blocks (LCBs) longer than 100bp and found in all genomes were extracted with Mauve’s stripSubsetLCBs program, aligned with Mafft ^86^ and concatenated into host mitochondrial and bacterial core genomes. Phylogenetic trees were produced from these core genomes using the GTR model and 100 bootstraps in PhyML-3.1 ^87^.

We compared host and symbiont evolutionary rates by estimating the divergence at synonymous sites for each host pair. Using the Biopython toolkit ^88^, we extracted the nucleic and amino acid sequences of 13 conserved mitochondrial and 718 bacterial core protein-coding genes (see below). Amino acid sequences were then aligned with Muscle ^89^ and reverse translated into codon alignments using the “build” function from the Biopython codonalign package. The mitochondrial and bacterial codon-based alignments were then each concatenated into two genome-wide alignments with complete (no gaps, no N) lengths of 10417 bp and 662118 bp, respectively. We assessed substitution saturation by plotting transitions and transversions against adjusted genetic distance. Pairwise synonymous (dS) substitution rates were computed using the Maximum-Likelihood method ^90^ implemented in the Biopython codonalign package. The source code was slightly modified to accommodate for ambiguous bases in the mitochondrial genomes.

### Identification of bacterial core genes

Because of low structural differences among genomes, orthologous genes could be inferred based on homology and position ^91^. A list of positional homologs with a minimum identity of 30% and a minimum coverage of 60% was exported from the Mauve alignments. Additional maps with a stricter identity criterion (60% identity, 80% coverage) were produced from the alignments of multiple subsets of symbiont genomes. The consensus of these orthologous maps yielded 749 core genes (Figure S2, Table S1) including 718 core protein-coding genes ranging from 138 bp to 4554 bp (average 975bp) (Table S3).

### Bayesian concordance analyses

We used Bucky v.1.2 ^92^ to estimate the proportion of core protein-coding genes supporting each topology. Putative recombination breakpoints within the 718 core protein-coding genes previously found were identified with GARD and the KH test ^93,94^. Using a false positive discovery rate threshold of 5%, recombination was found in 66 genes which were thus split into multiple contiguous non-recombining gene segments at the inferred breakpoints prior to phylogenetic inference.

Bucky takes as input the posterior distribution of topologies for each gene (or gene segment). These distributions were each obtained from 800 trees generated in MrBayes v.3.2.7a ^95^ using a Gamma + I rate variation across sites. These trees represented a well-mixed sample of the tree space after convergence of four independent Markov Chain Monte Carlo (MCMC) chains which were each run for 2,000,000 generations after an initial 100,000 generations burn-in period. Trees were sampled every 10,000 generations to avoid autocorrelation. Parameter optimization for the MCMCs was performed by assessing convergence and mixing of both continuous parameters of the model and tree topologies using the R package RWTY v.1.0.2 ^96^.

In Bucky, two independent MCMC runs were carried out using the prior assumption that all genes shared the same topology (alpha=0). MCMC runs performed 1,000,000 updates after an initial 10% burn-in period. One cold and three heated chains (swapping frequency =10) were used to improve mixing and convergence of all of the MCMC runs.

### Relaxed and positive selection detection

Relaxation of the strength of selection was detected in the symbiont genomes by two independent methods. First we use the Codon Deviation Coefficient (CDC) ^97^ to quantify codon usage bias on all protein-coding genes (Table S2) because this index does not require a priori knowledge of gene expression and is not biased by GC content. Second, we used RELAX ^49^ on individual core genes. RELAX detects change in the strength of selection between two groups by observing change in the distribution of ω (dN/dS ratio) classes in a branch-site random effects likelihood (BS-REL) framework between a set of test and reference branches. We compared *Ca*. Vesicomyosocius, *Ca*. Ruthia, and both clades together to the group composed of the free-living lineages.

To reduce false positives in phylogenetic selection tests ^98^, genes with significant evidence of recombination (see Bayesian concordance analyses) were excluded from these analyses. Episodic diversifying selection on individual lineages was identified on the remaining non-recombining 652 protein-coding genes using the adaptive Branch-site Random Effects Likelihood method (aBSRel ^99^). The Holm-Bonferroni correction for multiple testing was applied and threshold for detection was set to 10% false positive discovery rate. We used the hypergeometric test function dmvhyper from extraDistr v.1.8.11 ^100^ to test whether the genes under relaxed or positive selection represented a random subsample of all core genes according to SEED categories ^83^. The Fisher test ^101^ was applied to find SEED categories that were over-represented in the genes under relaxed or positive selection.

## Data availability

Symbiont genomes and Sequence Read Archives (SRAs) are available at the National Center for Biotechnology Information (NCBI) under the BioProject PRJNA641445. The mitochondrial genomes were deposited in GenBank under the references MT947381-MT947391.

Genome alignment files and Rmarkdown scripts of downstream analyses are available at https://github.com/maepz/VesicSymb_Evolution

## Authors contributions

CRY designed the study; CRY and YJW contributed to data collection; MP, CB, and CRY performed analysis, BA contributed to data interpretation. And all authors co-wrote the manuscript.

---

**Table S6.**
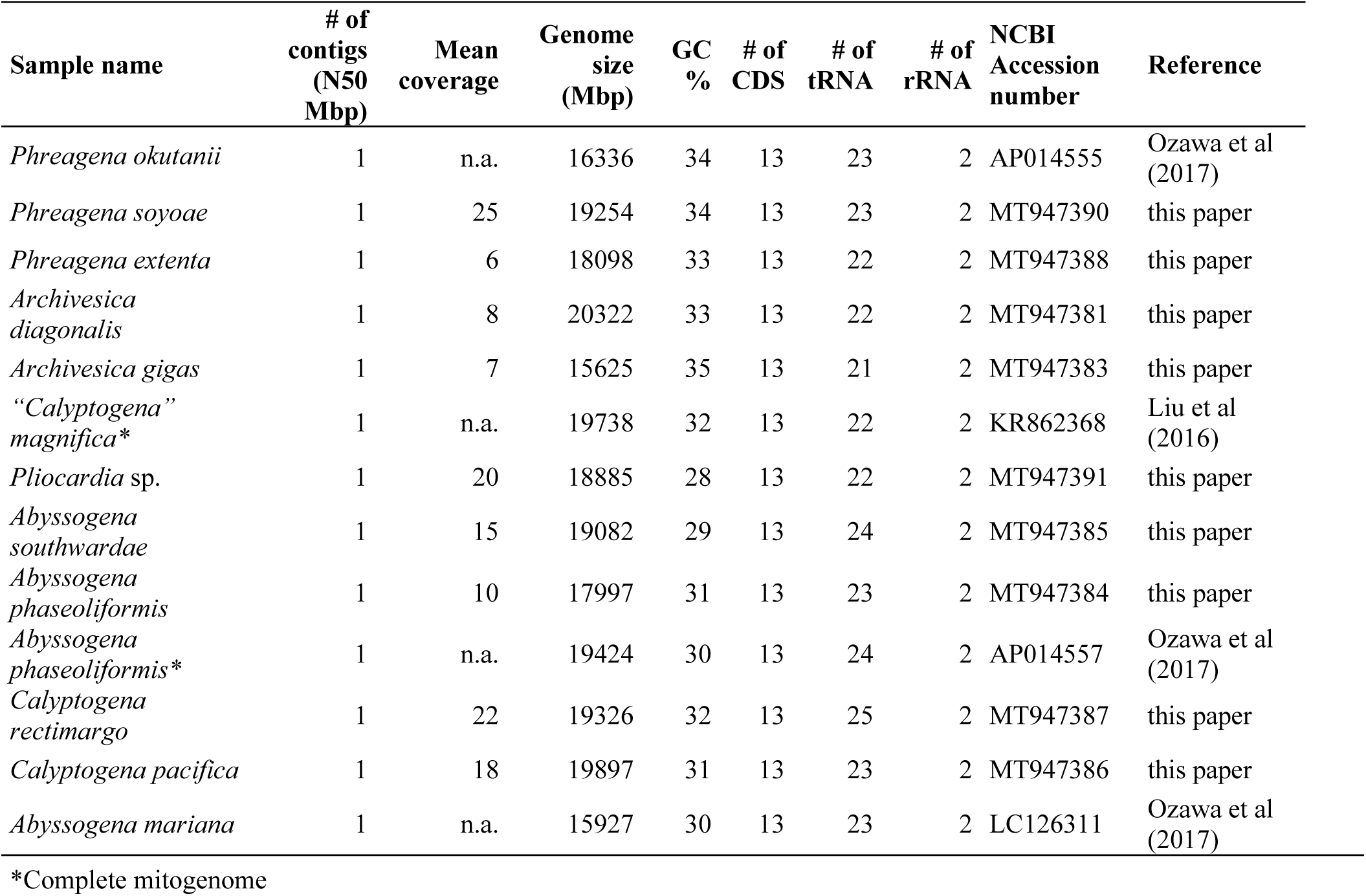
Assembly and annotation statistics for mitochondrial genomes in this study.

## Figures and captions

**Figure S1.**
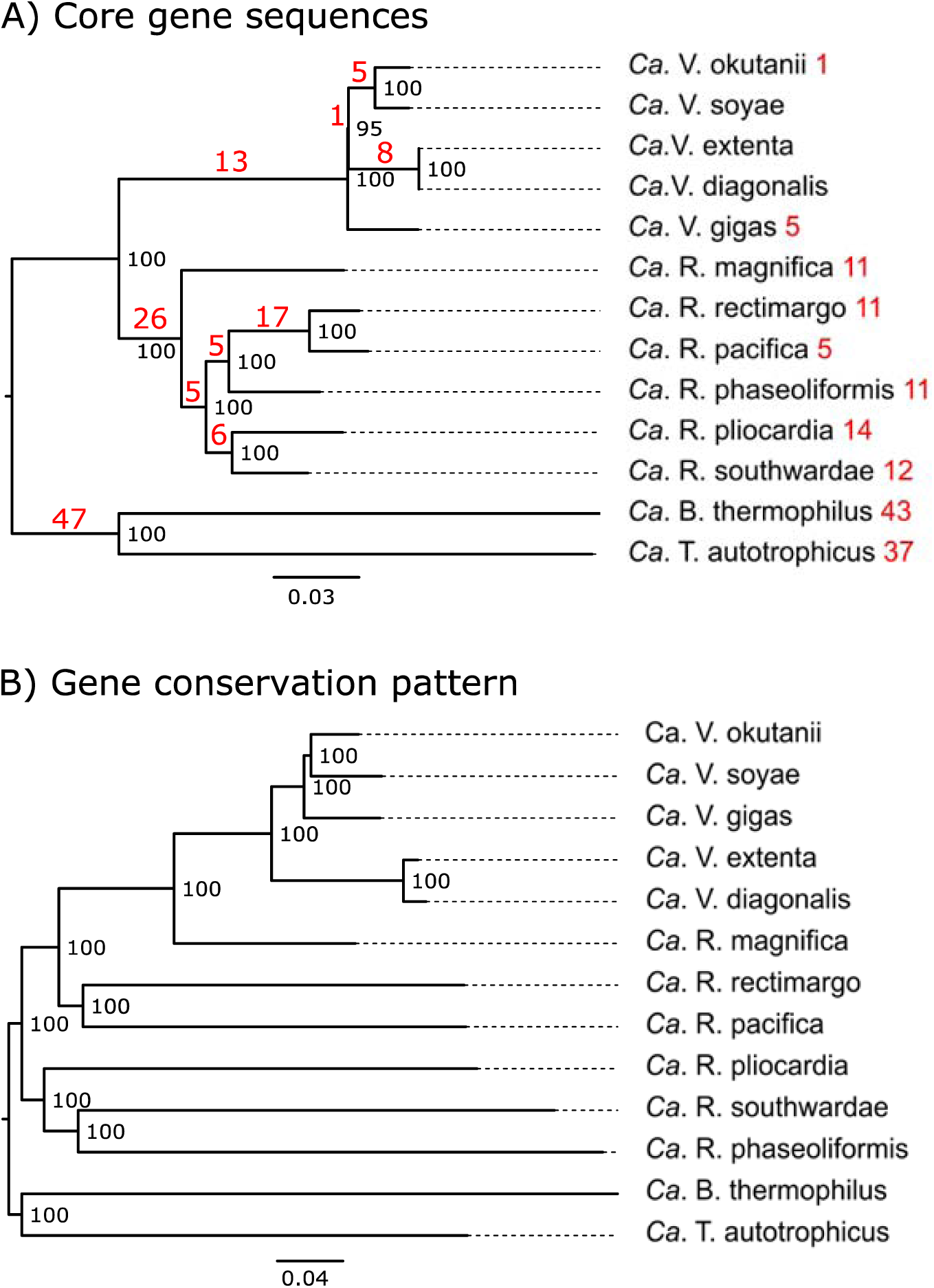
Distance-based neighbour-joining trees established from A) a concatenated alignment of 652 non-recombining core protein-coding genes sequences (618342 bp, HKY nucleotide substitution model). In red are the number of genes under episodic diversifying selection in each branch; B) the presence/absence of positionally orthologous genes (Jaccard distance on 6110 genes). Numbers above branches are bootstrap support values.

**Figure S2.**
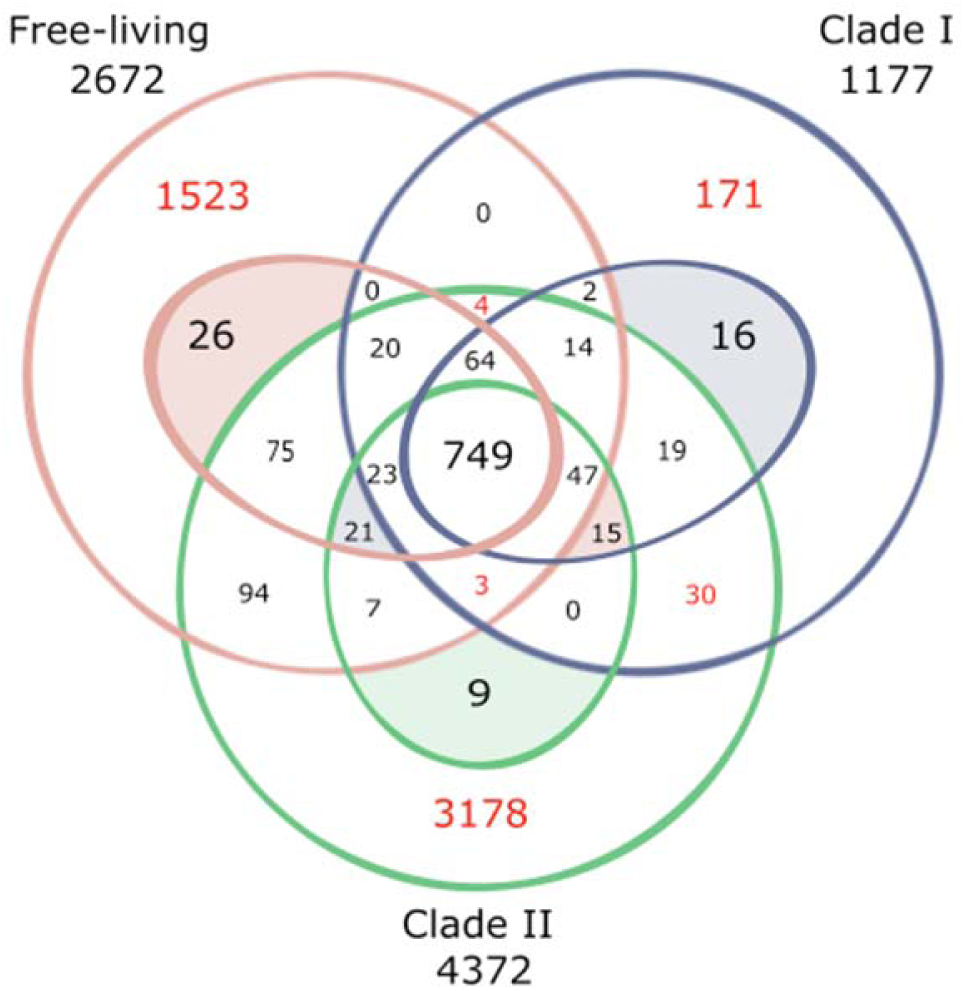
Venn diagram representing the 6110 unique and shared putative genes amongst the free-living, *Ca*. Vesicomyosocius and *Ca*. Ruthia. The outer circles represent the pan-genome while the inner circles represent the core-genome of the groups. Free-living: *Ca*. B. thermophilus and *Ca*. T. autotrophicus); *Ca*. Ruthia: *Ca*. R. magnifica, *Ca*. R. phaseoliformis, *Ca*. R. pacifica, *Ca*. R. rectiomargo, *Ca*. R. pliocardia, and *Ca*. R. southwardae; *Ca*. Vesicomyosocius: *Ca*. V. okutanii, *Ca*. V. soyoae, *Ca*. V. diagonalis-extenta, and *Ca*. V. gigas. Groups in which more than 50% of the genes are unannotated are identified in red. The complete orthology is available in Table S1.

**Figure S3.**
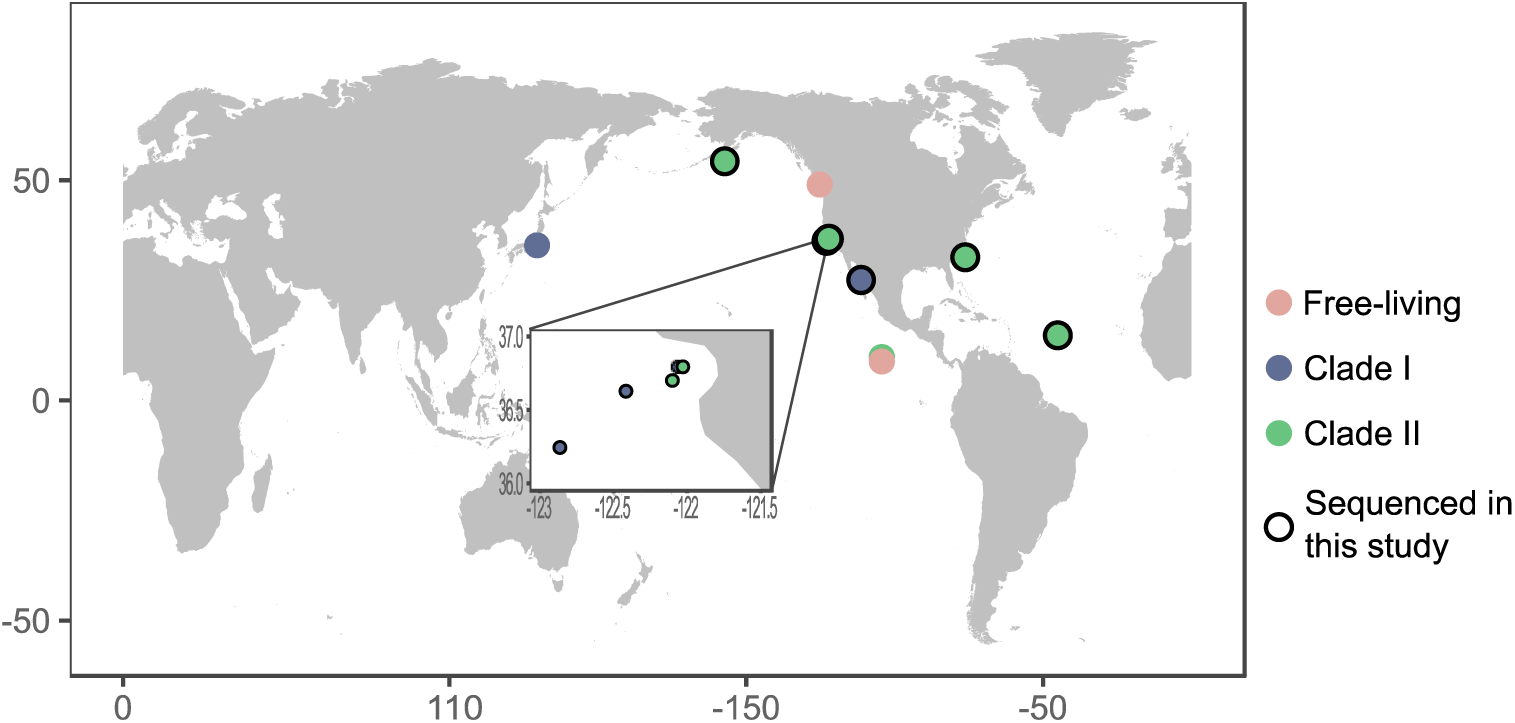
Sampling locations. Inset depicts samples collected from varying depths in Monterey Bay.

## References

1. Roger, A. J., Muñoz-Gómez, S. A. & Kamikawa, R. The Origin and Diversification of Mitochondria. Current Biology 27, R1177–R1192 (2017).

2. McFadden, G. I. Origin and Evolution of Plastids and Photosynthesis in Eukaryotes. Cold Spring Harb Perspect Biol 6, a016105 (2014).

3. Cavanaugh, C. M. Microbial Symbiosis: Patterns of Diversity in the Marine Environment. American Zoologist 34, 79–89 (1994).

4. Clavijo, J. M., Donath, A., Serôdio, J. & Christa, G. Polymorphic adaptations in metazoans to establish and maintain photosymbioses. Biological Reviews 93, 2006–2020 (2018).

5. Garate, L., Sureda, J., Agell, G. & Uriz, M. J. Endosymbiotic calcifying bacteria across sponge species and oceans. Scientific Reports 7, 43674 (2017).

6. Bright, M. & Bulgheresi, S. A complex journey: transmission of microbial symbionts. Nat Rev Micro 8, 218–230 (2010).

7. Baumann, P. et al. Genetics, physiology, and evolutionary relationships of the genus Buchnera: intracellular symbionts of aphids. Annu. Rev. Microbiol. 49, 55–94 (1995).

8. Wernegreen, J. J. Endosymbiont evolution: predictions from theory and surprises from genomes. Annals of the New York Academy of Sciences 1360, 16–35 (2015).

9. Fisher, R. M., Henry, L. M., Cornwallis, C. K., Kiers, E. T. & West, S. A. The evolution of host-symbiont dependence. Nature Communications 8, 15973 (2017).

10. Husnik, F. & Keeling, P. J. The fate of obligate endosymbionts: reduction, integration, or extinction. Current Opinion in Genetics & Development 58–59, 1–8 (2019).

11. Kuwahara, H. et al. Reductive genome evolution in chemoautotrophic intracellular symbionts of deep-sea Calyptogena clams. Extremophiles 12, 365–374 (2008).

12. Itoh, T., Martin, W. & Nei, M. Acceleration of genomic evolution caused by enhanced mutation rate in endocellular symbionts. PNAS 99, 12944–12948 (2002).

13. Moran, N. A., McLaughlin, H. J. & Sorek, R. The Dynamics and Time Scale of Ongoing Genomic Erosion in Symbiotic Bacteria. Science 323, 379–382 (2009).

14. Mendonça, A. G., Alves, R. J. & Pereira-Leal, J. B. Loss of Genetic Redundancy in Reductive Genome Evolution. PLOS Computational Biology 7, e1001082 (2011).

15. Lhee, D. et al. Evolutionary dynamics of the chromatophore genome in three photosynthetic Paulinella species. Scientific Reports 9, 2560 (2019).

16. Wernegreen, J. J. & Moran, N. A. Evidence for genetic drift in endosymbionts (Buchnera): analyses of protein-coding genes. Mol Biol Evol 16, 83–97 (1999).

17. Moran, N. A. Accelerated evolution and Muller’s rachet in endosymbiotic bacteria. PNAS 93, 2873–2878 (1996).

18. Kuo, C.-H., Moran, N. A. & Ochman, H. The consequences of genetic drift for bacterial genome complexity. Genome Res. 19, 1450–1454 (2009).

19. Muller, H. J. The relation of recombination to mutational advance. Mutation Research/Fundamental and Molecular Mechanisms of Mutagenesis 1, 2–9 (1964).

20. Bennett, G. M. & Moran, N. A. Heritable symbiosis: The advantages and perils of an evolutionary rabbit hole. PNAS 112, 10169–10176 (2015).

21. Andersson, S. G. E. & Kurland, C. G. Reductive evolution of resident genomes. Trends in Microbiology 6, 263–268 (1998).

22. Andersson, J. O. & Andersson, S. G. Genome degradation is an ongoing process in Rickettsia. Mol Biol Evol 16, 1178–1191 (1999).

23. Peek, A. S., Vrijenhoek, R. C. & Gaut, B. S. Accelerated evolutionary rate in sulfur-oxidizing endosymbiotic bacteria associated with the mode of symbiont transmission. Mol Biol Evol 15, 1514–1523 (1998).

24. Cary, S. C. & Giovannoni, S. J. Transovarial inheritance of endosymbiotic bacteria in clams inhabiting deep-sea hydrothermal vents and cold seeps. PNAS 90, 5695–5699 (1993).

25. Ikuta, T. et al. Surfing the vegetal pole in a small population: extracellular vertical transmission of an ‘intracellular’ deep-sea clam symbiont. Royal Society Open Science 3, 160130 (2016).

26. Peek, A. S., Feldman, R. A., Lutz, R. A. & Vrijenhoek, R. C. Cospeciation of chemoautotrophic bacteria and deep sea clams. PNAS 95, 9962–9966 (1998).

27. Stewart, F. J., Young, C. R. & Cavanaugh, C. M. Evidence for homologous recombination in intracellular chemosynthetic clam symbionts. Mol. Biol. Evol. 26, 1391–1404 (2009).

28. Peek, A. S., Gustafson, R. G., Lutz, R. A. & Vrijenhoek, R. C. Evolutionary relationships of deep-sea hydrothermal vent and cold-water seep clams (Bivalvia: Vesicomyidae): results from the mitochondrial cytochrome oxidase subunit I. Marine Biology 130, 151–161 (1997).

29. Moran, N. A., Munson Mark A., Baumann Paul & Ishikawa Hajime. A molecular clock in endosymbiotic bacteria is calibrated using the insect hosts. Proceedings of the Royal Society of London. Series B: Biological Sciences 253, 167–171 (1993).

30. Ferri, E. et al. New Insights into the Evolution of Wolbachia Infections in Filarial Nematodes Inferred from a Large Range of Screened Species. PLOS ONE 6, e20843 (2011).

31. Johnson, S. B., Krylova, E. M., Audzijonyte, A., Sahling, H. & Vrijenhoek, R. C. Phylogeny and origins of chemosynthetic vesicomyid clams. Systematics and Biodiversity 15, 346–360 (2017).

32. Krylova, E. M. & Sahling, H. Vesicomyidae (Bivalvia): Current Taxonomy and Distribution. PLoS ONE 5, e9957 (2010).

33. Audzijonyte, A., Krylova, E. M., Sahling, H. & Vrijenhoek, R. C. Molecular taxonomy reveals broad trans-oceanic distributions and high species diversity of deep-sea clams (Bivalvia: Vesicomyidae: Pliocardiinae) in chemosynthetic environments. Systematics and Biodiversity 10, 403–415 (2012).

34. MolluscaBase. MolluscaBase. Vesicomyidae Dall & Simpson, 1901. http://www.marinespecies.org/aphia.php?p=taxdetails&id=23140#distributions (2019).

35. Newton, I. L. G. et al. The Calyptogena magnifica Chemoautotrophic Symbiont Genome. Science 315, 998–1000 (2007).

36. Kuwahara, H. et al. Reduced Genome of the Thioautotrophic Intracellular Symbiont in a Deep-Sea Clam, Calyptogena okutanii. Current Biology 17, 881–886 (2007).

37. Kuwahara, H. et al. Loss of genes for DNA recombination and repair in the reductive genome evolution of thioautotrophic symbionts of Calyptogena clams. BMC Evolutionary Biology 11, 285 (2011).

38. Tamas, I. et al. 50 Million Years of Genomic Stasis in Endosymbiotic Bacteria. Science 296, 2376–2379 (2002).

39. Roeselers, G. et al. Complete genome sequence of Candidatus Ruthia magnifica. Stand Genomic Sci 3, 163–173 (2010).

40. Anantharaman, K., Breier, J. A., Sheik, C. S. & Dick, G. J. Evidence for hydrogen oxidation and metabolic plasticity in widespread deep-sea sulfur-oxidizing bacteria. PNAS 110, 330–335 (2013).

41. Liu, H., Cai, S., Zhang, H. & Vrijenhoek, R. C. Complete mitochondrial genome of hydrothermal vent clam Calyptogena magnifica. Mitochondrial DNA Part A 27, 4333–4335 (2016).

42. Ozawa, G. et al. Updated mitochondrial phylogeny of Pteriomorph and Heterodont Bivalvia, including deep-sea chemosymbiotic Bathymodiolus mussels, vesicomyid clams and the thyasirid clam Conchocele cf. bisecta. Mar Genomics 31, 43–52 (2017).

43. Yang, M., Gong, L., Sui, J. & Li, X. The complete mitochondrial genome of Calyptogena marissinica (Heterodonta: Veneroida: Vesicomyidae): Insight into the deep-sea adaptive evolution of vesicomyids. PLoS One 14, (2019).

44. Ip, J. C.-H. et al. Host-Endosymbiont Genome Integration in a Deep-Sea Chemosymbiotic Clam. Molecular Biology and Evolution (2020) doi:10.1093/molbev/msaa241.

45. Stackebrandt, E. & Goebel, B. M. Taxonomic Note: A Place for DNA-DNA Reassociation and 16S rRNA Sequence Analysis in the Present Species Definition in Bacteriology. International Journal of Systematic and Evolutionary Microbiology, 44, 846–849 (1994).

46. Cohan, F. M. Sexual Isolation and Speciation in Bacteria. Genetica 116, 359–370 (2002).

47. Fujiwara, Y. et al. Phylogenetic characterization of endosymbionts in three hydrothermal vent mussels: influence on host distributions. Marine Ecology Progress Series 208, 147–155 (2000).

48. Petersen, J. M. et al. Hydrogen is an energy source for hydrothermal vent symbioses. Nature 476, 176–180 (2011).

49. Wertheim, J. O., Murrell, B., Smith, M. D., Kosakovsky Pond, S. L. & Scheffler, K. RELAX: Detecting Relaxed Selection in a Phylogenetic Framework. Mol Biol Evol 32, 820–832 (2015).

50. Burke, G. R. & Moran, N. A. Massive Genomic Decay in Serratia symbiotica, a Recently Evolved Symbiont of Aphids. Genome Biol Evol 3, 195–208 (2011).

51. Stewart, F. J., Young, C. R. & Cavanaugh, C. M. Lateral Symbiont Acquisition in a Maternally Transmitted Chemosynthetic Clam Endosymbiosis. Mol Biol Evol 25, 673–687 (2008).

52. Decker, C., Olu, K., Arnaud-Haond, S. & Duperron, S. Physical Proximity May Promote Lateral Acquisition of Bacterial Symbionts in Vesicomyid Clams. PLoS One 8, (2013).

53. Ozawa, G. et al. Ancient occasional host switching of maternally transmitted bacterial symbionts of chemosynthetic vesicomyid clams. Genome biology and evolution 9, 2226–2236 (2017).

54. Kojima, S. The distribution and the phylogenies of the species of genus Calyptogena and those of vestimentiferans around Japan. JAMSTEC Journal of Deep-Sea Research 11, 243–248 (1995).

55. Goffredi, S. K. & Barry, J. P. Species-specific variation in sulfide physiology between closely related Vesicomyid clams. Marine Ecology Progress Series 225, 227–238 (2002).

56. Breusing, C., Johnson, S. B., Vrijenhoek, R. C. & Young, C. R. Host hybridization as a potential mechanism of lateral symbiont transfer in deep-sea vesicomyid clams. Molecular Ecology 28, 4697–4708 (2019).

57. Naito, M. & Pawlowska, T. E. Defying Muller’s Ratchet: Ancient Heritable Endobacteria Escape Extinction through Retention of Recombination and Genome Plasticity. mBio 7, e02057–15 (2016).

58. Pérez-Brocal, V. et al. A small microbial genome: the end of a long symbiotic relationship? Science 314, 312–313 (2006).

59. Koga, R. & Moran, N. A. Swapping symbionts in spittlebugs: evolutionary replacement of a reduced genome symbiont. ISME J 8, 1237–1246 (2014).

60. Sudakaran, S., Kost, C. & Kaltenpoth, M. Symbiont Acquisition and Replacement as a Source of Ecological Innovation. Trends in Microbiology 25, 375–390 (2017).

61. Chong, R. A. & Moran, N. A. Evolutionary loss and replacement of Buchnera, the obligate endosymbiont of aphids. The ISME Journal 12, 898 (2018).

62. Latorre, A. & Manzano□Marín, A. Dissecting genome reduction and trait loss in insect endosymbionts. Annals of the New York Academy of Sciences 1389, 52–75 (2017).

63. Saki, M. & Prakash, A. DNA damage related crosstalk between the nucleus and mitochondria. Free Radical Biology and Medicine 107, 216–227 (2017).

64. Chu, H. & Mazmanian, S. K. Innate immune recognition of the microbiota promotes host-microbial symbiosis. Nature immunology 14, 668–675 (2013).

65. Oldroyd, G. E. Speak, friend, and enter: signalling systems that promote beneficial symbiotic associations in plants. Nature Reviews Microbiology 11, 252–263 (2013).

66. Brownlie, J. C., Adamski, M., Slatko, B. & McGraw, E. A. Diversifying selection and host adaptation in two endosymbiont genomes. BMC Evolutionary Biology 7, 68 (2007).

67. Dale, C. & Moran, N. A. Molecular Interactions between Bacterial Symbionts and Their Hosts. Cell 126, 453–465 (2006).

68. Chong, R. A., Park, H. & Moran, N. A. Genome Evolution of the Obligate Endosymbiont Buchnera aphidicola. Mol Biol Evol (2019) doi:10.1093/molbev/msz082.

69. Wagner, G. P. & Gabriel, W. Quantitative Variation in Finite Parthenogenetic Populations: What Stops Muller’s Ratchet in the Absence of Recombination? i>Evolution 44, 715–731 (1990).

70. Rand, D. M., Haney, R. A. & Fry, A. J. Cytonuclear coevolution: the genomics of cooperation. Trends in Ecology & Evolution 19, 645–653 (2004).

71. Fares, M. A., Moya, A. & Barrio, E. Adaptive evolution in GroEL from distantly related endosymbiotic bacteria of insects. Journal of Evolutionary Biology 18, 651–660 (2005).

72. Howe, D. K. & Denver, D. R. Muller’s Ratchet and compensatory mutation in Caenorhabditis briggsae mitochondrial genome evolution. BMC Evolutionary Biology 8, 62 (2008).

73. Castillo, D. M. & Pawlowska, T. E. Molecular Evolution in Bacterial Endosymbionts of Fungi. Mol Biol Evol 27, 622–636 (2010).

74. Ohishi, K. et al. Long-term Cultivation of the Deep-Sea Clam Calyptogena okutanii: Changes in the Abundance of Chemoautotrophic Symbiont, Elemental Sulfur, and Mucus. The Biological Bulletin 230, 257–267 (2016).

75. Andrews, S. et al. astQC. (2012).

76. Zerbino, D. R. & Birney, E. Velvet: algorithms for de novo short read assembly using de Bruijn graphs. Genome Res. 18, 821–829 (2008).

77. Kearse, M. et al. Geneious Basic: An integrated and extendable desktop software platform for the organization and analysis of sequence data. Bioinformatics 28, 1647–1649 (2012).

78. Langmead, B. & Salzberg, S. L. Fast gapped-read alignment with Bowtie 2. Nat Meth 9, 357–359 (2012).

79. Li, H. et al. The Sequence Alignment/Map format and SAMtools. Bioinformatics 25, 2078–2079 (2009).

80. Bankevich, A. et al. SPAdes: A New Genome Assembly Algorithm and Its Applications to Single-Cell Sequencing. Journal of Computational Biology 19, 455–477 (2012).

81. Hahn, C., Bachmann, L. & Chevreux, B. Reconstructing mitochondrial genomes directly from genomic next-generation sequencing reads—a baiting and iterative mapping approach. Nucleic Acids Res 41, e129–e129 (2013).

82. Tillich, M. et al. GeSeq – versatile and accurate annotation of organelle genomes. Nucleic Acids Res 45, W6–W11 (2017).

83. Overbeek, R. et al. The SEED and the Rapid Annotation of microbial genomes using Subsystems Technology (RAST). Nucleic Acids Res 42, D206–D214 (2014).

84. Darling, A. E., Mau, B. & Perna, N. T. progressiveMauve: Multiple Genome Alignment with Gene Gain, Loss and Rearrangement. PLoS ONE 5, e11147 (2010).

85. Tesler, G. GRIMM: genome rearrangements web server. Bioinformatics 18, 492–493 (2002).

86. Katoh, K. & Standley, D. M. MAFFT Multiple Sequence Alignment Software Version 7: Improvements in Performance and Usability. Mol Biol Evol 30, 772–780 (2013).

87. Guindon, S. et al. New Algorithms and Methods to Estimate Maximum-Likelihood Phylogenies: Assessing the Performance of PhyML 3.0. Syst Biol 59, 307–321 (2010).

88. Cock, P. J. A. et al. Biopython: freely available Python tools for computational molecular biology and bioinformatics. Bioinformatics 25, 1422–1423 (2009).

89. Edgar, R. C. MUSCLE: multiple sequence alignment with high accuracy and high throughput. Nucleic Acids Res. 32, 1792–1797 (2004).

90. Goldman, N. & Yang, Z. A codon-based model of nucleotide substitution for protein-coding DNA sequences. Mol Biol Evol 11, 725–736 (1994).

91. Lemoine, F., Lespinet, O. & Labedan, B. Assessing the evolutionary rate of positional orthologous genes in prokaryotes using synteny data. BMC Evol Biol 7, 237 (2007).

92. Larget, B. R., Kotha, S. K., Dewey, C. N. & Ané, C. BUCKy: Gene tree/species tree reconciliation with Bayesian concordance analysis. Bioinformatics 26, 2910–2911 (2010).

93. Kishino, H. & Hasegawa, M. Evaluation of the maximum likelihood estimate of the evolutionary tree topologies from DNA sequence data, and the branching order in hominoidea. J Mol Evol 29, 170–179 (1989).

94. Kosakovsky Pond, S. L., Posada, D., Gravenor, M. B., Woelk, C. H. & Frost, S. D. W. GARD: a genetic algorithm for recombination detection. Bioinformatics 22, 3096–3098 (2006).

95. Ronquist, F. et al. MrBayes 3.2: Efficient Bayesian Phylogenetic Inference and Model Choice Across a Large Model Space. Systematic Biology 61, 539–542 (2012).

96. Warren, D. L., Geneva, A. J. & Lanfear, R. RWTY (R We There Yet): An R package for examining convergence of Bayesian phylogenetic analyses. Molecular Biology and Evolution msw279 (2017) doi:10.1093/molbev/msw279.

97. Zhang, Z. et al. Codon Deviation Coefficient: a novel measure for estimating codon usage bias and its statistical significance. BMC Bioinformatics 13, 43 (2012).

98. Pond, S. L. K., Frost, S. D. W. & Muse, S. V. HyPhy: hypothesis testing using phylogenies. Bioinformatics 21, 676–679 (2005).

99. Smith, M. D. et al. Less Is More: An Adaptive Branch-Site Random Effects Model for Efficient Detection of Episodic Diversifying Selection. Mol Biol Evol 32, 1342–1353 (2015).

100. Wołodźko, T. extraDistr: Additional Univariate and Multivariate Distributions. (2019).

101. Fisher, S. R. A. Confidence Limits for a Cross-Product Ratio. Australian Journal of Statistics 4, 41–41 (1962).

